# A KASP Marker for the Potato Late Blight Resistance Gene *RB*/*Rpi-blb1*

**DOI:** 10.1101/2023.02.22.529539

**Authors:** Peyton L. Sorensen, Grace Christensen, Hari S. Karki, Jeffrey B. Endelman

## Abstract

The disease late blight is a threat to potato production worldwide, making genetic resistance an important target for breeding. The resistance gene *RB*/*Rpi-blb1* is effective against most strains of the causal pathogen, *Phytophthora infestans*. Until now, potato breeders have utilized a Sequence Characterized Amplified Region (SCAR) marker to screen for *RB*. Our objective was to design and validate a Kompetitive Allele Specific PCR (KASP) marker, which has advantages for high-throughput screening. First, the accuracy of the SCAR marker was confirmed in two segregating tetraploid populations. Then, using whole genome sequencing data for two *RB*-positive segregants and a diverse set of 23 *RB*-negative varieties, a SNP in the 5’ untranslated (UTR) region was identified as unique to *RB*. The KASP marker based on this SNP, which had 100% accuracy in the cultivated diversity panel, was used to generate diploid breeding lines containing *RB*. The KASP marker is publicly available for others to utilize.

## INTRODUCTION

Late blight is caused by the oomycete pathogen *Phytophthora infestans* (Mont. De Bary), which spreads rapidly under moist conditions and causes significant economic injury to potato (*S. tuberosum*) crops worldwide (Judelson and Blanco 2005; Fry 2008). The annual global cost for late blight, due to both crop injury and fungicide use to prevent the disease, has been estimated at more than US$ 5 billion (Haverkort et al. 2008). Hence, there are strong economic and environmental incentives to breed varieties with genetic resistance. Unfortunately, *P. infestans* has a history of overcoming R genes, particularly those introgressed from the wild potato relative *S. demissum* (Fry 2008).

The gene *RB*/*Rpi-blb1* from *S. bulbocastanum* has a NLR (nucleotide binding site and leucine-rich repeat) motif and exhibits broad-spectrum resistance to most strains of *P. infestans* (Song et al. 2003; van der Vossen et al. 2003; Chen et al. 2012; Haverkort et al. 2016). Because *S. bulbocastanum* is not sexually compatible with *S. tuberosum*, *RB* was introduced through somatic hybridization several decades ago (Helgeson et al. 1998). To enable marker-assisted breeding, Colton et al. (2006) developed a sequence characterized amplified region (SCAR) marker, which uses two primers to generate an amplicon through polymerase chain reaction (PCR). While this marker was reported to be effective, its throughput is limited by the need to assay the amplicon-size polymorphism.

The advent of KASP (Kompetitive Allele Specific PCR) markers has allowed breeders to screen for single nucleotide polymorphisms (SNP) more efficiently. The KASP system is based on two allele-specific primers, each with a corresponding fluorophore, and a common reverse primer. Based on the fluorescence ratio for the two alleles, all three genotypes in a diploid are readily distinguished, and the different heterozygote dosages in a polyploid can sometimes be inferred (Uitdewilligen et al. 2013; Kante et al. 2021).

The main objective of this research was to design and validate a KASP marker for the *RB* gene. We also illustrate the power of new software tools for haplotype analysis in outbred F1 populations and describe the generation of *S. tuberosum* diploids containing *RB*.

## MATERIALS & METHODS

### Plant Material

*RB*-positive clones were obtained from multiple sources. The somatic hybrids J101, J103, J117, and J136 were obtained from D. Halterman (USDA-ARS, Madison, WI), as well as K41, (full name J101K6A6K41) which is the BC_3_ offspring between J101 and *S. tuberosum* varieties Katahdin (K) and Atlantic (A). Two other *RB*-positive clones derived from backcrossing, Sbu8.5 and J138K6A22, were obtained from G. Porter (University of Maine) and D. Douches (Michigan State University), respectively. In 2019 K41 was used as the female parent in crosses with two *RB*-negative clones, W13NYP102-7 and W6609-3, which created the populations W19003B (n=21) and W19004B (n=12), respectively.

### Experimental Protocols

Greenhouse experiments were conducted with 16 h light/8 h dark and an average day temperature of 70 °F (21 °C) and night temperature of 60 °F (15.5 °C). Detached leaf assays (DLA) were conducted on each offspring from the W19003B and W19004B populations (one pot per offspring) when plants were 7 weeks old. One leaf was collected from the middle third of each plant and placed in a tray with wet paper towels. The inoculum of the *P. infestans* isolate US-23 was adjusted to 50,000 zoospores per 1 mL of water, and then 4-6 10 μL droplets were placed on the abaxial side of each leaflet (Karki and Halterman 2021). Visual inspection 5 days after inoculation was used to classify samples as susceptible or resistant.

Some changes to the published protocol for the 1+1’ SCAR marker (Colton et al. 2006) were needed to obtain satisfactory results. Primers were obtained from IDT (Coralville, Iowa), and DNA was extracted from leaf tissue using the Qiagen DNeasy plant kit (Hilden, Germany). A 25 μL PCR reaction was conducted using 12.5 μL Promega 2x PCR Master Mix, 9.5 μL of nuclease free water (NFW), 1 μL each of the 10 μM 1+1’ forward and reverse primers, and 1 μL of template DNA diluted to 60-120 ng/μL. The PCR protocol was run on a Bio-Rad C1000 thermal cycler (Hercules, CA) as follows: 95 °C for 7 minutes; 37 cycles of 95 °C for 20 seconds, 50 °C for 20 seconds, and 72 °C for 2 minutes; 72 °C for 7 minutes. A 1.2% agarose gel was made with 1x TBE and 5 μL of SYBR^®^ Safe DNA gel stain. A loading dye mix was made with two-parts Promega Blue/Orange 6x loading dye and one-part 6x MassRuler loading dye. 2.5 μL of the loading dye mix was combined with 10 μL of the 1+1’ PCR product and loaded into the gel. The gel was electrophoresed with the VWR^®^ Real Time Electrophoresis System (Radnor, Pennsylvania) in a 0.5x TBE bath for 90 minutes at 50 V. The banding results from the gel were visualized with a blue LED transilluminator.

The KASP marker was designed by LGC Biosearch Technologies (Teddington, UK) based on the target SNP (G/A at 42,613 bp) and flanking sequence in the reference genome (Genbank ID AY303170). The Project ID for ordering the primers is 2571.006, with Assay ID RB_086. The assay was conducted in a 10 μL reaction volume, using 5 μL of 2x KASP master mix, 0.14 μL of assay mix, and 5 μL of template DNA (diluted to 20-50 ng/μL). Thermal cycling used the standard 61–55°C touchdown protocol: 94 °C for 15 minutes; 10 cycles of 94 °C for 20 seconds and 61 °C for 1 minute with a decrease of 0.6 °C every cycle; 26 cycles of 94 °C for 20 seconds and 55 °C for 1 minute; a final plate reading step at 30 °C for 1 minute. Clusters were defined based on the slope of their regression line according to the ratio of HEX: FAM fluorescence. If the clusters were not easily distinguishable then a recycling protocol was run as follows: 3 cycles of 94 °C for 20 seconds and 57 °C for 60 seconds; a final plate reading step at 30 °C for 60 seconds.

### Data Analysis

Genetic linkage mapping was conducted using marker data (File S1) from the potato v3 SNP array (Felcher et al. 2012; Vos et al. 2015); the genotyping service provider was Neogen (Lincoln, NE). Genotype calls were made using the package fitPoly (Voorrips et al. 2011; Zych et al. 2019) in R (R Core Team 2021). Parental phasing and haplotype reconstruction of the W19003B and W19004B populations were performed using PolyOrigin (Zheng et al. 2021) in Julia v1.7.2. Mapping of binary trait loci (BTL) for the SCAR marker and detached leaf assay results (File S2) was conducted using R package diaQTL (Amadeu et al. 2021).

Bioinformatic analysis was based on the reference genome AY303170, which corresponds to the minus strand for *RB*. Illumina 2×150 bp reads for W19003B-1 and W19003B-4 were aligned using BWA-MEM version 0.7.17 (Li 2013). SAMtools version 1.13 (Danecek et al. 2021) was used to remove unmapped reads and PCR duplicates. A VCF file was generated using FreeBayes v1.3.5 (Garrison and Marth 2012) and processed using the vcfR package (Kraus and Grünwald 2017). Sequencing data from Hardigan et al. (2017) for 23 *RB*-negative *S. tuberosum* varieties was downloaded from the European Nucleotide Archive (accession PRJNA378971) and processed as above.

## RESULTS

### SCAR Marker Validation

We created two tetraploid F1 populations (W19003B and W19004B) segregating for *RB*, which shared a common resistant parent K41. 17 of 33 progeny were classified as resistant based on a detached leaf assay, which is consistent with the 1:1 ratio expected for a single copy of *RB* in the resistant parent. 20 of 33 progeny tested positive for the SCAR marker, and the two-way table revealed 7 discrepancies between the two assays (Table 1).

**Table 1.**
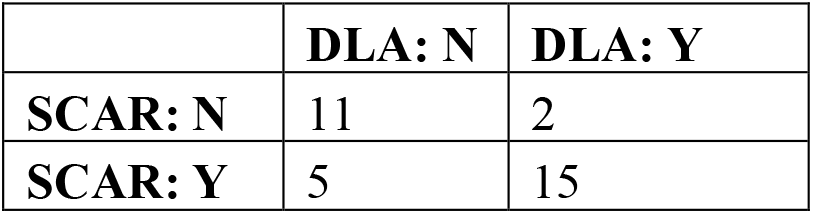
Two-way table for the presence (Y) vs. absence (N) of the *RB* gene in 33 F1 progeny, as determined by detached leaf assay (DLA) or SCAR marker.

To identify which method was more reliable for detecting the presence of *RB*, the populations were SNP array genotyped to construct linkage maps and perform binary trait locus (BTL) mapping. Both traits localized to chromosome 8, which is the known location of *RB*, but the SCAR peak was higher (Fig. 1). 100% of the variance for the SCAR marker was explained at the BTL peak, indicating perfect agreement with transmission of the parental haplotypes. The haplotype effect plot at the BTL peak (Fig. 2) shows the SCAR marker was specific to the second haplotype (numbering is arbitrary) of the resistant parent, designated K41.2. By contrast, only 65% of the variance for the detached leaf assay could be explained, which we attributed to micro-environmental or genetic background effects (Śliwka et al. 2012).

**Figure 1.**
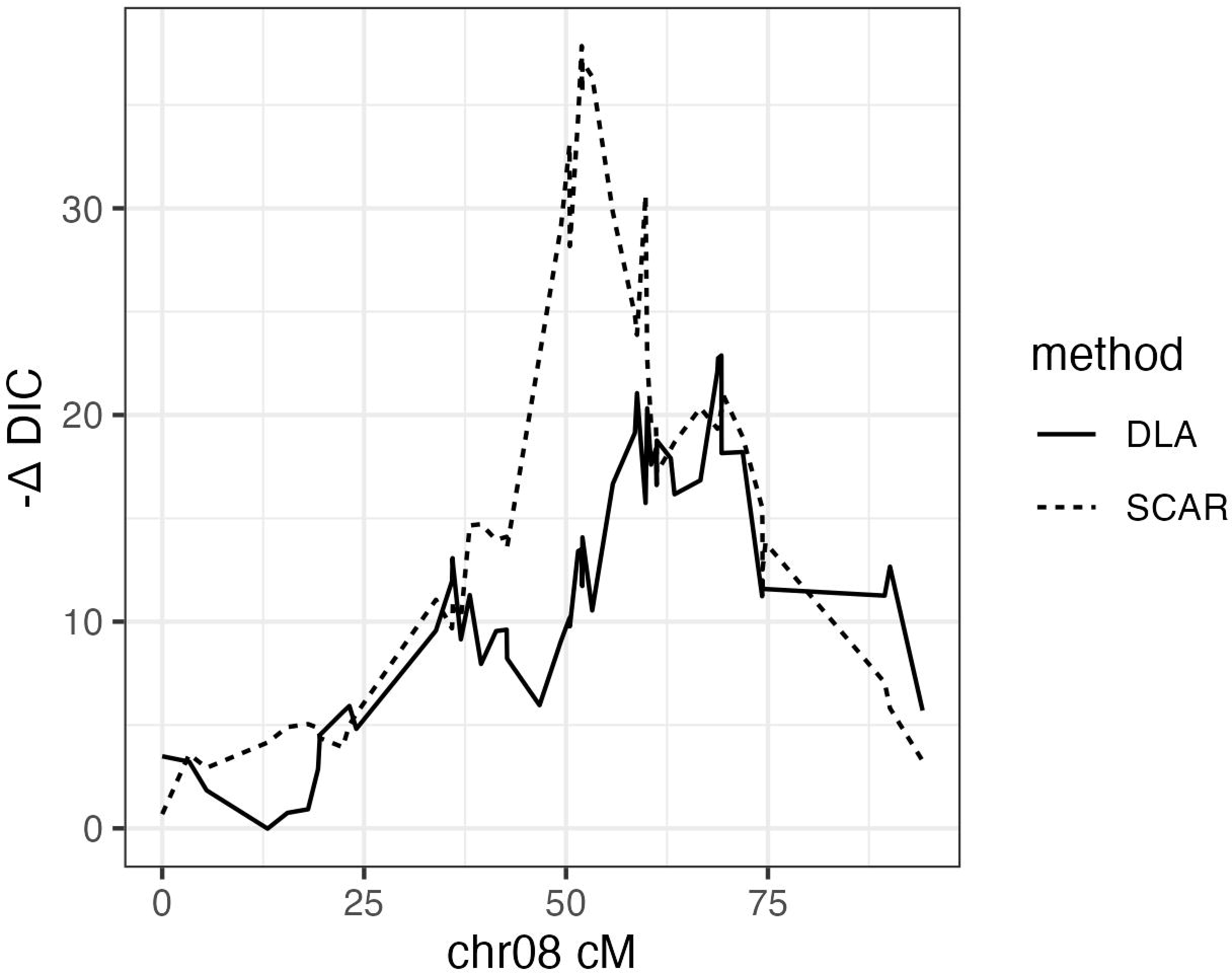
Linkage mapping in a tetraploid multi-parental population, using binary response data from a detached leaf assay (DLA) vs. a SCAR marker for the *RB* gene.

**Figure 2.**
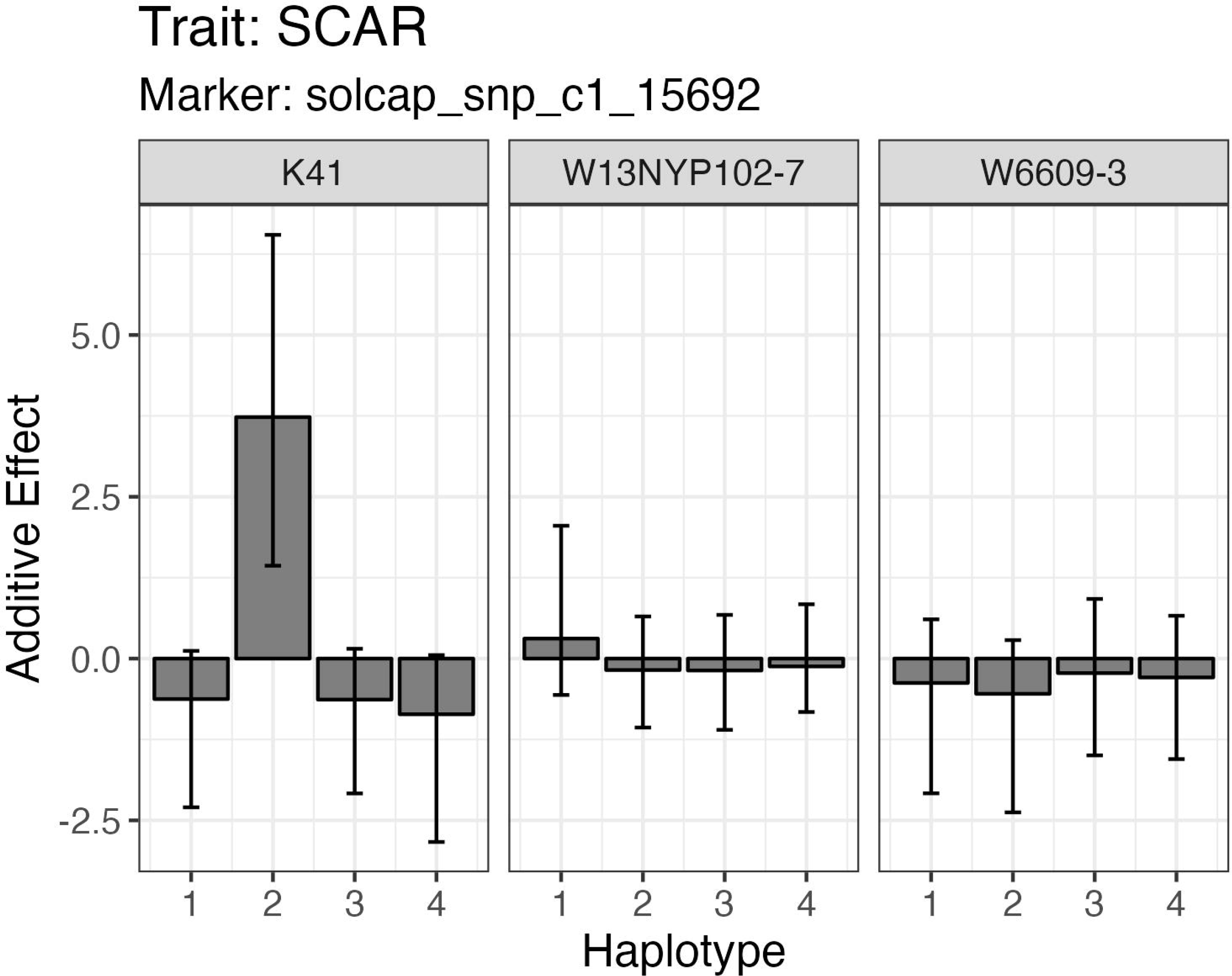
Additive effects of the four haplotypes for each parent of the multi-parental population. The large positive effect for haplotype K41.2 indicates the presence of *RB*.

### KASP Marker

For KASP marker design, our goal was to identify a SNP unique to the *RB* gene. Two *RB*-positive individuals from the W19003B population—progeny 1 and 4—were sequenced with 2×150 bp Illumina reads. Based on sequencing yields of 89 and 75 Gb, respectively, and the monoploid potato genome size of 840 Mb, the depth of coverage was estimated at 90–100X. Variant discovery was performed after aligning reads to the 79 kb reference sequence AY303170, which corresponds to the minus strand of *RB* and contains other R gene analogs. From the alignment depth (Fig. 3), we deduced that paralogous reads were being mapped to the two exons, which would confound attempts to identify a haplotype-specific SNP. Three SNPs in the intron and one in the 5’ UTR were identified as candidates based on the simplex (i.e., single copy) dosage of the reference (*RB*) allele. Using whole-genome sequencing data for 23 tetraploid varieties presumed to be *RB*-negative (Hardigan et al. 2017), only two of the four candidate SNPs (A/T at 41,673 and G/A at 42,613 bp) were bi-allelic and homozygous for the alternate allele in all 23 clones.

**Figure 3.**
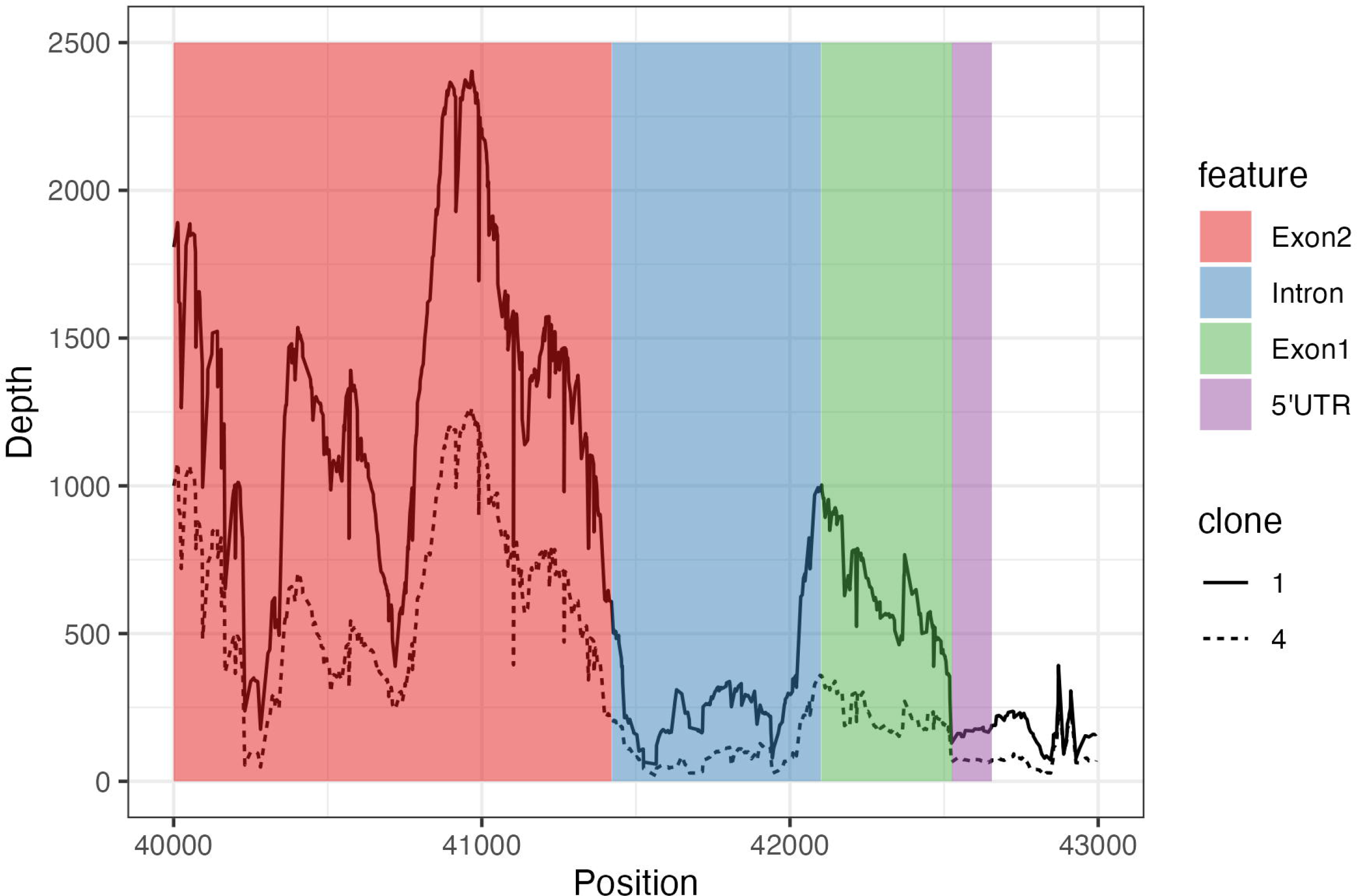
Read depth from whole-genome sequencing of two *RB*-positive progeny (1 and 4) of the tetraploid W19003B population. Based on the expected coverage of 90–100X, paralogous reads were aligned in the exons.

The SNP at 42,613 bp, which is 86 bp upstream of the start codon, was selected for KASP marker design. The validation panel included five of the original *S. bulbocastanum* hybrids (J101, J103, J117, J136, J138) and/or their descendants (File S3). *RB*-negative samples spanned all four of the main cultivated groups in N. America (chip, russet, red, yellow). There were no discrepancies between the KASP and SCAR marker results (Fig. 4).

**Figure 4.**
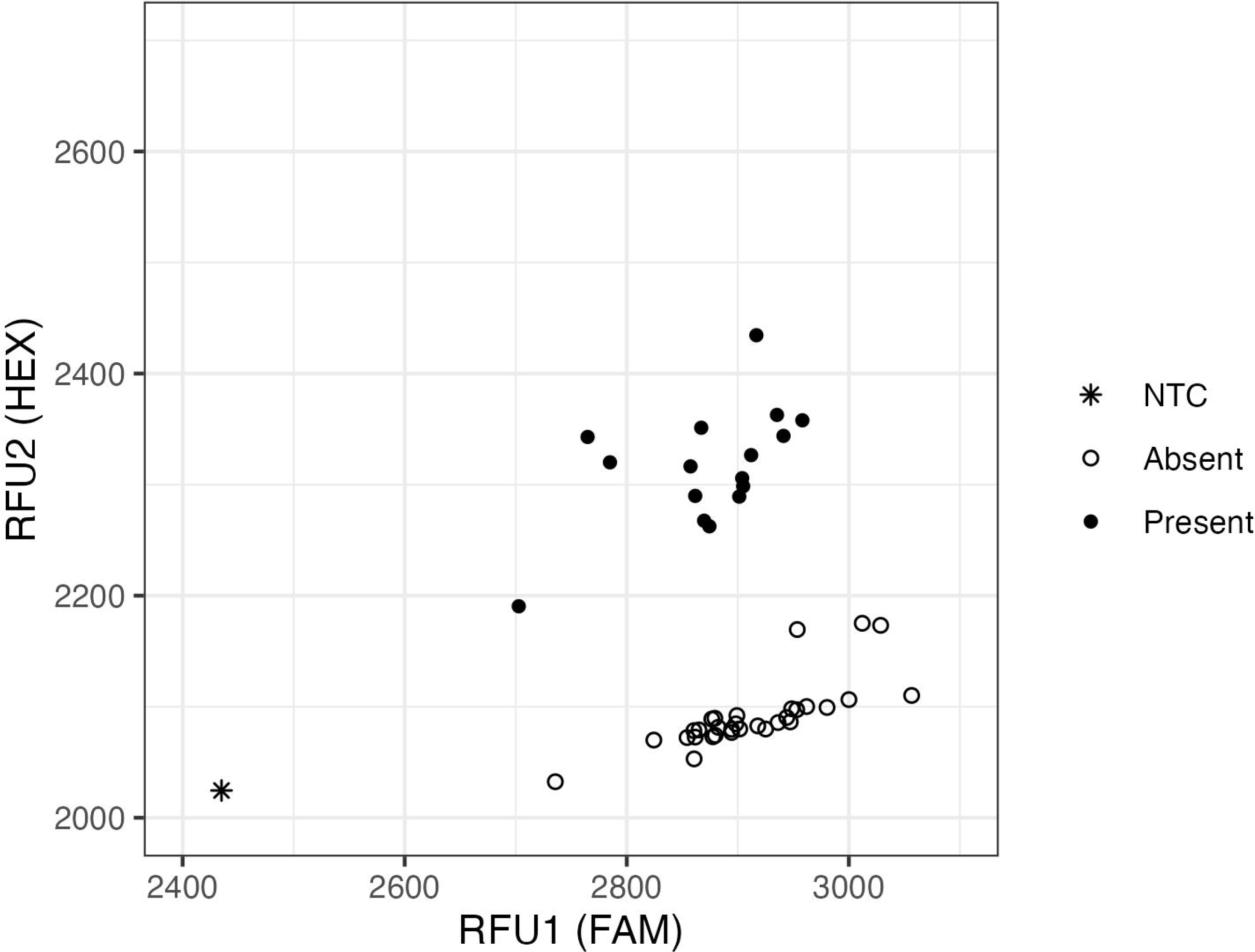
Results with the KASP marker were in perfect agreement with the SCAR marker (present/absent) in a diverse set of cultivated N. American germplasm. NTC = no template control.

### Diploid Breeding

*RB*-positive individuals from the W19003B population were selected as targets for haploid induction with the pollinator IVP101 (Hermsen and Verdenius 1973). Four individuals produced over 20 seeds without the embryo spot (Table 2), which indicates a putative dihaploid. 28 progeny were classified as diploid based on chloroplast counts in the stomatal guard cells (Ordoñez et al. 2017), of which 14 tested positive for *RB* using the KASP marker, which matches the expected 1:1 segregation ratio. One *RB*-positive dihaploid, W19003B-12-DH13, was selected for breeding based on its female fertility when pollinated with W2×082-(14/20)-13, an F_2_ clone homozygous for *Sli* (Song and Endelman 2022). *RB* was successfully transmitted to the F1 progeny, and three *RB*-positive individuals were selected for further breeding based on the yield of tubers and true seeds (Table 3).

**Table 2.** Haploid induction results for *RB*-positive clones from the W19003B population.

| **Tetraploid Mother** | **Seeds** | **Seeds without embryo spot** | **Putative dihaploids** | **Dihaploids with *RB*** |
| --- | --- | --- | --- | --- |
| W19003B-4 | 209 | 94 | 7 | 2 |
| W19003B-6 | 45 | 24 | 6 | 2 |
| W19003B-12 | 135 | 91 | 14 | 10 |
| W19003B-21 | 46 | 23 | 1 | 0 |
| **Totals** | 435 | 232 | 28 | 14 |

**Table 3.**
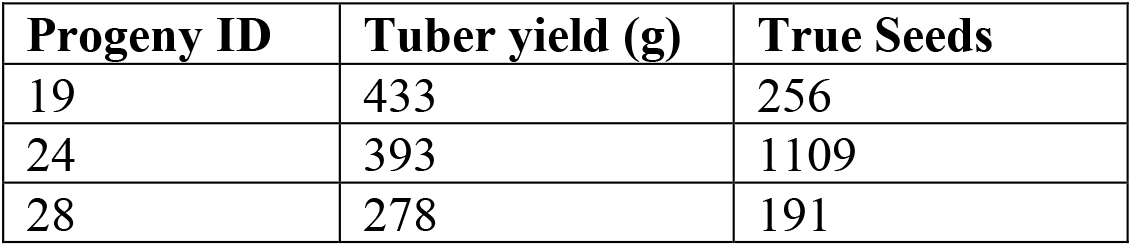
Selected *RB*-positive diploids from the F1 population W2x150 = W19003B-12-DH13 × W2x082-(14/20)-13. Yields based on one plant in a 3.78L pot. True seeds based on pollination with W2x082-(14/20)-13.

## DISCUSSION

It has been 20 years since *RB*/*Rpi-blb1* was cloned (Song et al. 2003; van der Vossen et al. 2003) and over 15 since the publication of the SCAR marker (Colton et al. 2006), but potato breeders have yet to commercialize a variety with this gene for the N. American market without genetic engineering (Brown-Donovan et al. 2021). (There has been greater success, and arguably greater effort, in the European market [Rakosy-Tican et al. 2020].) One contributing factor may be that, even after multiple rounds of backcrossing to *S. tuberosum*, *RB*-positive clones derived from somatic hybrids can be poorly adapted to modern production systems. For example, the *RB*-donor in this study (K41) has excessive stolons. Ideally, marker-assisted backcrossing should be used to eliminate undesirable *S. bulbocastanum* haplotypes while retaining *RB*. But tetrasomic inheritance makes this process inherently slow, and until recently there was not convenient software to select on parental haplotypes in tetraploid F1 populations (Zheng et al. 2021). Marker-assisted backcrossing at the diploid level is more efficient (Su et al. 2020), and the results of this study make us optimistic that inbred clones homozygous for *RB* will be available soon.

As in other crops, KASP markers have significantly increased the efficiency of breeding for pest resistance in potato. Some programs have the capacity to run these assays in-house, particularly when the number of samples is limited. For screening many hundreds or thousands of individuals, the KASP genotyping service offered through the CGIAR Excellence in Breeding (EiB) initiative has been valuable for public breeding programs. The first potato KASP markers developed for the EiB service targeted variants linked to *Ry_adg_* (Herrera et al. 2018; Kante et al. 2021) and *Ry_sto_* (Nie et al. 2016), which confer extreme resistance to potato virus Y. More recent targets include the late blight resistance genes *R2/Rpi-abpt/Rpi-blb3* on chr04 (Meade et al. 2020; Karki et al. 2021) and *R8/Rpi-smira2* on chr09 (Kante et al. 2021). According to Haverkort et al. (2016), *R8/Rpi-smira2* compares favorably to *RB*/*Rpi-blb1* in terms of durability, while *R2/Rpi-abpt/Rpi-blb3* only offers “intermediate” spectrum resistance.

Good stewardship requires the pursuit of varieties with multiple R genes for late blight. Accomplishing this in tetraploids through sexual hybridization is difficult because late blight R genes are still at low frequency in elite N. American germplasm, requiring positive selection for multiple R genes in addition to agronomic and quality traits. These challenges have made genetic engineering an attractive alternative (Zhu et al. 2012; Ghislain et al. 2019), and pyramiding multiple R genes is also more straightforward in diploids. Our KASP marker for *RB* provides yet another tool for achieving the long-standing goal of durable resistance against late blight.

## Supplemental Material

File S1. SNP array marker data for the W19003B and W19004B populations.

File S2. Phenotype data for the detached leaf assay (DLA) and SCAR marker.

File S3. Validation panel for the KASP marker.

## Author Contributions

PS: Formal analysis, Investigation, Writing – original draft, Writing – review & editing. GC: Methodology, Writing – review & editing. HK: Investigation, Writing – review & editing. JBE: Conceptualization, Formal analysis, Funding acquisition, Supervision, Writing – review & editing.

## Acknowledgments

Financial support was provided by the USDA National Institute of Food Agriculture Award 2019-51181-30021.

## Data Availability

SNP array and phenotype data are provided as Supplemental Files S1 and S2, respectively. Whole-genome sequence data for W19003B-1 and W19003B-4 is available from the NCBI Sequence Read Archive (SRA) under BioProject ID PRJNA934950.

